# Somatic *TYK2* activating mutations in tumor-infiltrating T cells promote anti-cancer immunity

**DOI:** 10.1101/2025.10.13.682217

**Authors:** Zhijie Li, Meng-Hsiung Hsieh, Alisa Arutyunova, John Evans, Tao Liu, Andrew R.J. Lawson, Pantelis A. Nicola, Gerda Kildisiute, Heather E. Machado, Ahmad Al Kawam, Rui Wang, Qiyu Zeng, Nisi Jiang, Xun Wang, Gregory Mannino, Xiongzhao Ren, Xiangyi Fang, Tripti Sharma, Suman Komjeti, David Hsiehchen, Scott Hayton, Simon Brunner, Iñigo Martincorena, Peter J. Campbell, Hao Zhu

## Abstract

Cancer cells evolve to increase fitness and evade the immune system, but it is not clear if tumor infiltrating lymphocytes (TILs) undergo selection for somatic mutations that augment anti-cancer immunity. Using single molecule whole exome sequencing in TILs, we identified somatic mutations in Tyrosine-protein kinase 2 (*TYK2*), several of which increased TYK2 phosphorylation, JAK-STAT signaling, and interferon gamma signaling. Among these mutations, *TYK2^D810V^* was recurrently observed in patients with diverse cancer types. In vitro, *TYK2^D810V^* enhanced T cells effector functions and cytokine production. We generated mice with a germline *Tyk2^D807V^* mutation that recapitulated the human *TYK2^D810V^*mutation. These mice were healthy and did not develop autoimmune disorders. However, they possessed enhanced anti-tumor effects in the context of syngeneic and autochthonous cancer models. Adoptive therapy with *Tyk2^D807V^* CD8⁺ T cells also decreased cancer growth. Thus, naturally occurring mutations in non-malignant lymphocytes can antagonize cancer and inspire new immunotherapies strategies.

## Background

Somatic mutations are widespread in normal tissues prior to the development of cancer. The hematopoietic system was the first tissue in which a large array of somatic mutations were identified, a phenomenon called clonal hematopoiesis of indeterminate potential (CHIP). CHIP is the prototypical paradigm for how somatic mutations impact human health, but most hematopoietic mutations appear to have maladaptive consequences ^1–5^. These mutations (*DNMT3A*, *TET2, ASXL1*) are predictive of leukemia risk and are associated with increased inflammation in cardiac, neurologic, and liver diseases ^2,6–10^. Interestingly, hematopoietic mutations found in CHIP have been associated with accelerated solid tumor development, but surprisingly have also been associated with altered immune checkpoint inhibitor (ICI) responses and may enhance engineered cellular therapies ^11–14^. More recently, somatic mutations identified in lymphomas such as the *CARD11-PIK3R3* fusion were found to augment anti-cancer immune function when introduced into CAR-T cells ^15^. Somatic loss of function mutations in *TNFAIP3*, which are frequently found in B cell lymphomas, were also found to have anticancer immune effects when identified in TILs ^16^. A unifying feature of these observations is that the immunity augmenting mutations in *DNMT3A*, *TET2*, *ASXL1, CARD11-PIK3R3*, and *TNFAIP3* are all commonly associated with leukemia/lymphoma development. A knowledge gap in the field is whether there are naturally occurring, non-malignancy associated somatic mutations in normal immune cells that can increase anti-cancer immunity.

To further explore this question, we performed single molecule somatic genomic sequencing of tumor infiltrating lymphocytes (TILs) from cancer patients. Among several genes, we identified multiple mutations in *TYK2*, one of which appeared recurrently in three patients with distinct tumor types. TYK2 was the first described member of the JAK family, and plays a pivotal role in inflammatory signalling ^17–19^. In immune cells, TYK2 has been implicated as a signaling component downstream of IFN-α, IL-6, IL-10, and IL-12 ^20–22^. Hypomorphic *TYK2* germline mutations have been linked to a decreased risk of developing autoimmune diseases such as psoriasis ^23–26^, and deucravacitinib is a small-molecule TYK2 inhibitor that has been approved for the treatment of plaque psoriasis, highlighting the essential role of TYK2 in regulating immune signaling ^27–29^. Here, we asked if naturally occurring *TYK2* point mutations in T cells are sufficient to protect against cancer development or progression, and to uncover the precise mechanisms by which *TYK2* mutations may enhance anti-tumor effects. Our findings lay the foundation for identifying and utilizing somatic mutations as therapeutic concepts in immunotherapy. Identifying naturally occurring somatic mutations in non-malignant lymphocytes could lead to drug targets with reduced long-term potential for malignant transformation.

## Results

### TILs harbored somatic mutations in *TYK2*

To identify somatic mutations, we used whole-exome NanoSeq, a single-molecule duplex sequencing approach with error rates <5 errors per billion sites in single molecules of DNA that permits the sensitive detection of mutations with low variant allele frequencies (VAFs) within a heterogeneous population of cells ^30^. We applied this high-fidelity sequencing method to 153 TIL samples isolated from 46 patients with non-small cell lung cancer, melanoma, colorectal cancer, and renal cell carcinoma (**Fig. 1a,b**). The TIL compartment included purified populations of CD4⁺ T cells, CD8⁺ T cells, B cells, and NK cells. This analysis uncovered somatic mutations in several genes associated with T cell signaling and immune responses (see **Table S1**). The dnds algorithm was used to quantify positive selection at nucleotide resolution ^31^. We focused our analysis on *TYK2*, a key signaling kinase in the JAK-STAT pathway, and identified 54 somatic mutations across different TIL subsets (**Fig. 1c**). TYK2 belongs to the JAK family, which has four members (JAK1-3 and TYK2). Each protein is composed of seven homology domains called JH domains that include kinase (JH1) and pseudokinase (JH2) domains. The JH2 pseudokinase domain does not bind to ATP, but regulates kinase function ^32^.

**Figure 1.**
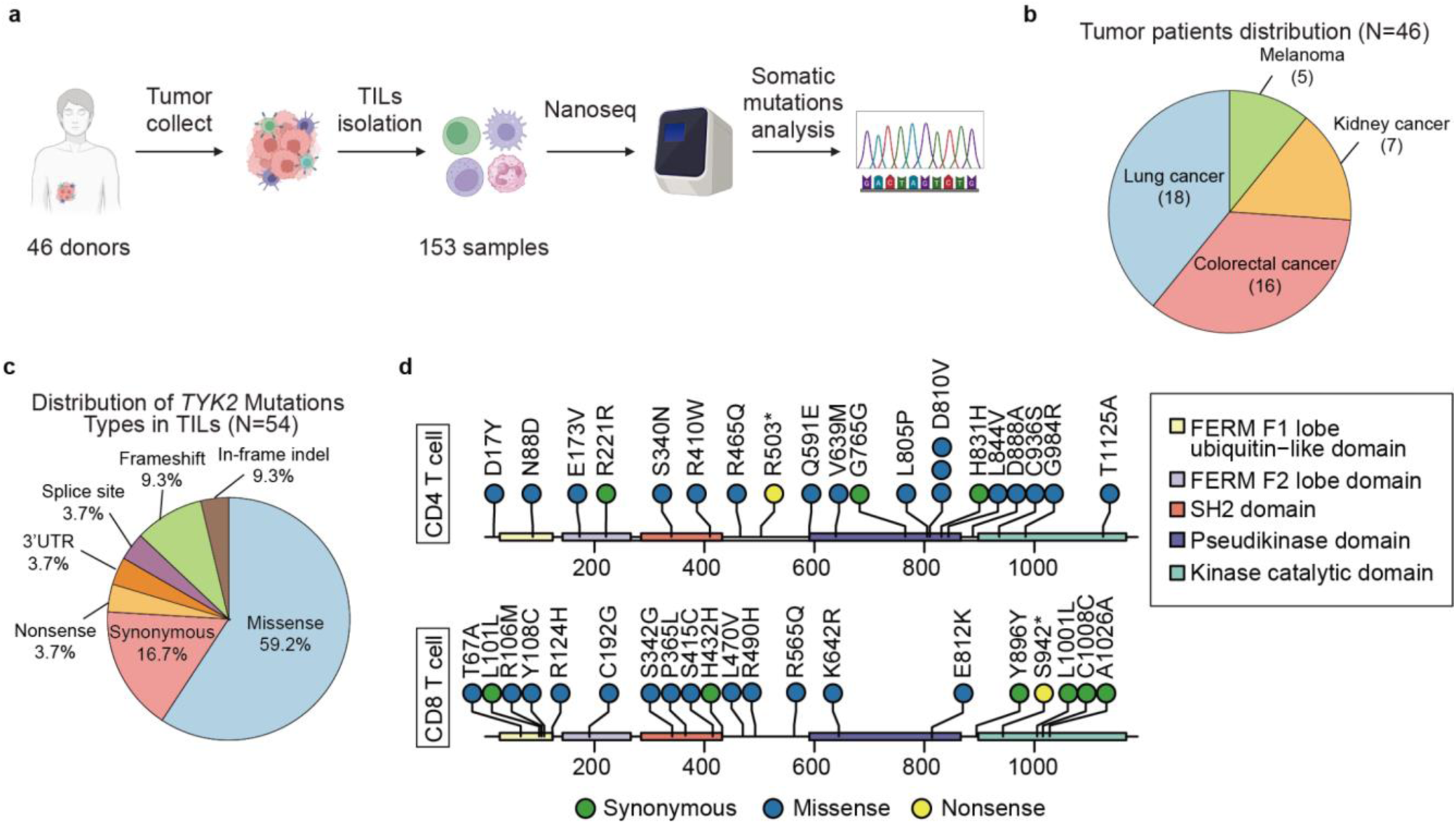
NanoSeq of TILs identified somatic mutations in *TYK2*. **a.** A schematic of somatic genomic sequencing of patient derived TILs. **b.** Distribution of tumor patients types. **c.** Classification of *TYK2* mutation types in TILs. **d.** Lollipop plots of TYK2 mutations in tumor-infiltrating CD4⁺ T cells and tumor-infiltrating CD8⁺ T cells.

Among the 54 *TYK2* mutations identified, missense mutations were the most common (59.2%), followed by synonymous (16.7%), in-frame indels (9.3%), frameshifts (9.3%), splice site (3.7%), nonsense (3.7%), and 3′UTR variants (3.7%) (**Fig. 1c**). Notably, we identified 15 missense *TYK2* mutations specifically in tumor-infiltrating CD4⁺ T cells, including three independent instances of *D810V* mutations in NSCLC, melanoma, and RCC patients (**Fig. 1d**). The remaining TYK2 mutations in other subsets were unique to individual patients, suggesting they may represent private events (**Fig. 1d** and Extended Data Fig. 1a). The recurrence of D810V raised the possibility that this alteration plays a shared role in multiple cancer types. Importantly, these mutations were not observed in the germline and are not common in CHIP.

### *TYK2^D810V^* enhanced T cells effector functions

We overexpressed each of the 15 *TYK2* missense mutants identified from CD4⁺ T cells in Jurkat T cells, an immortal human T cell line commonly used to study signaling. We reasoned that these mutations might modulate JAK-STAT signalling and impact T cell activation and effector functions. In comparison with wild-type (WT) controls, many of the point mutations altered JAK-STAT signaling. Four mutations (Q591E, L805P, D810V, C936S) increased TYK2 phosphorylation and downstream STAT1 and STAT3 activation vs. controls. Interestingly, D810V caused the most significant JAK-STAT activation (**Fig. 2a**). This suggested that many of the TYK2 somatic mutations could have a functional impact within TILs.

**Figure 2.**
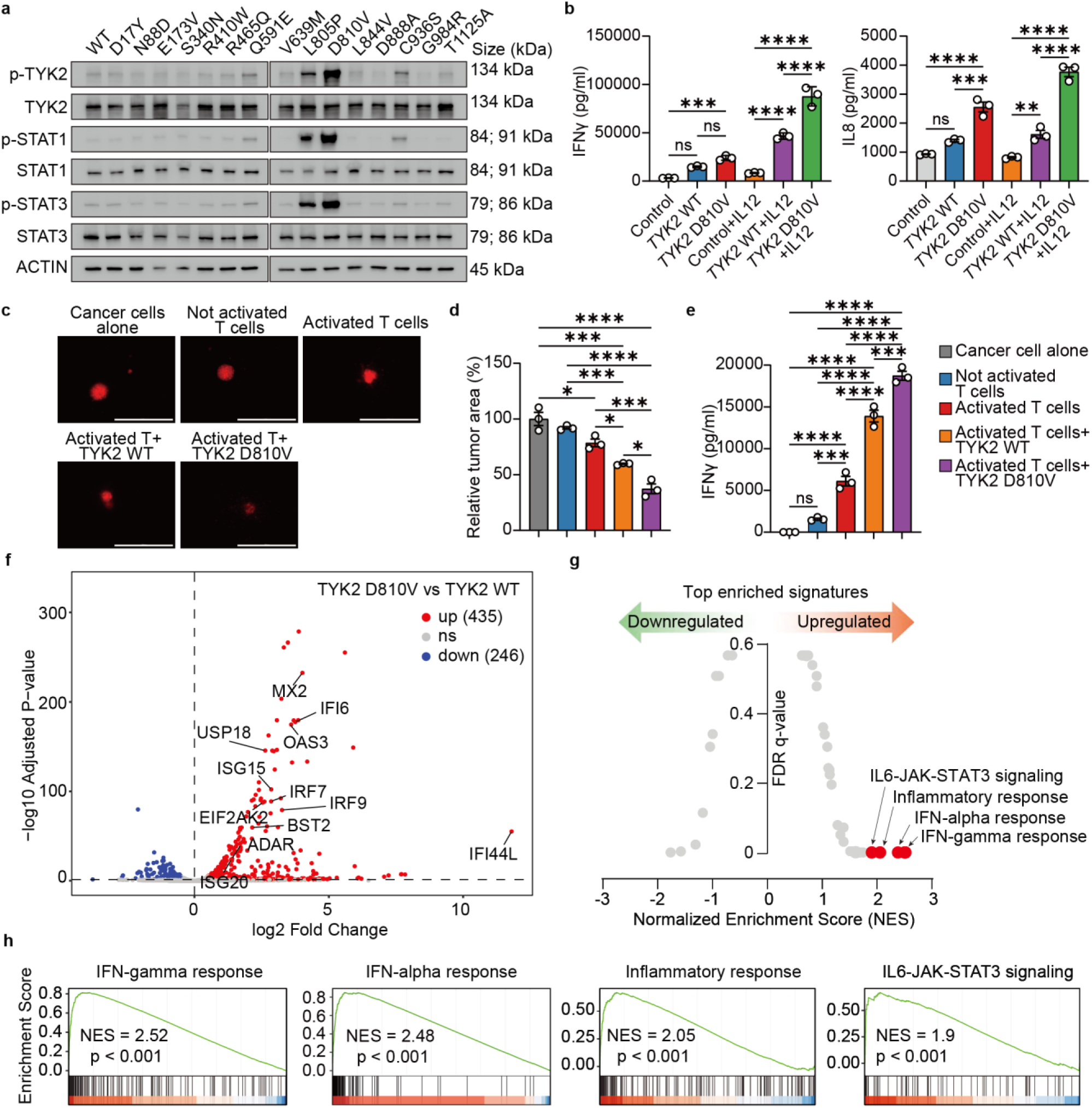
*TYK2^D810V^* expression enhances T cells effector function **a.** Western blot of Jurkat cells overexpressing 15 *TYK2* missense mutations. Representative blot from two independent experiments. **b.** IFNγ and IL8 levels measured in primary T cells expressing empty vector, *TYK2^WT^*, or *TYK2^D810V^* with or without IL12 (*n=*3 replicates per group). **c.** Representative images of A549 tumor co-cultured with primary T cells overexpressed empty vector, *TYK2^WT^*, or *TYK2^D810V^*, scale bars = 1000 μm (*n=*3 replicates per group). **d.** Relative tumor area of A549 tumor co-cultured with primary T cells overexpressed empty vector, *TYK2^WT^*, or *TYK2^D810V^*(*n*=3 replicates per group). **e.** IFNγ levels measured in A549 tumor cells co-cultured with primary T cells expressing empty vector, *TYK2^WT^*, or *TYK2^D810V^* (*n*=3 replicates per group). **f.** Volcano plot showing upregulated genes in Jurkat T cells overexpressing *TYK2^D810V^* compared to *TYK2^WT^* (*n=*3 replicates per group). **g.** Enriched gene signatures associated with up- or down-regulated genes associated with *TYK2^D810V^* compared to *TYK2^WT^*. **h.** GSEA showing upregulation of Interferon-gamma response, Interferon-alpha response, Interferon-gamma response and IL6-JAK-STAT3 signaling genes in *TYK2^D810V^* overexpressing Juakat T cells.

We focused on D810V, which is in the pseudokinase domain that regulates the kinase domain ^17^. To define the signaling changes downstream of *TYK2* mutations, we generated *TYK2* knockout 293T cells containing a STAT3 reporter. In these cells, overexpression of the *TYK2^D810V^* mutant resulted in higher STAT3 activation compared to overexpression of *TYK2^WT^* (Extended Data Fig. 2a). Expression of *TYK2^D810V^* in T cells also enhanced cytokine production, including IFN-γ and IL-8, compared to *TYK2^WT^* or control cells, and this effect was further potentiated by IL-12 stimulation, which stimulated the tyrosine phosphorylation of TYK2. This indicated that the *TYK2^D810V^* mutation enhanced both basal and cytokine-stimulated signaling responses (**Fig. 2b**). To further assess the impact of *TYK2^D810V^* on T cell mediated cytotoxicity, we performed a 3D tumor spheroid killing assay using A549 tumor cells co-cultured with T cells (Extended Data Fig. 2b). *TYK2^D810V^* expressing T cells exhibited enhanced tumor killing compared to controls (**Fig. 2c,d**). Culture media from *TYK2^D810V^* expressing T cells had increased IFN-γ, further supporting a gain-of-function phenotype associated with increased T cell effector activity (**Fig. 2e**).

To investigate the mechanisms by which *TYK2^D810V^* might influence inflammation, we performed RNA sequencing (RNA-seq) of human Jurkat T cells transduced with WT or *TYK2^D810V^*. 435 genes were upregulated with *TYK2^D810V^* overexpression, including known downstream targets of JAK-STAT (*MX2, IRF7, IRF9, OAS3, ISG15, ISG20, USP18, IFI6, IFI44L*) (**Fig. 2f**). Gene set enrichment analysis (GSEA) revealed enrichment of pathways associated with Interferon-gamma response, Interferon-alpha response, inflammatory responses, and JAK-STAT pathways, consistent with the known effects of TYK2 activation (**Fig. 2g-h**). This suggested that *TYK2^D810V^* drove a pro-inflammatory gene expression program by amplifying Interferon (IFN) responses and JAK–STAT signaling. For loss of function analysis, we performed single-cell RNA sequencing on primary human T cells with and without CRISPR-mediated *TYK2* deletion. This resulted in the downregulation of JAK–STAT and interferon signaling pathway genes in both CD4⁺ and CD8⁺ T cell subsets (Extended Data Fig. 2c-f). These transcriptional changes mirrored the reciprocal pattern observed in Jurkat T cells overexpressing *TYK2^D810V^*, reinforcing TYK2’s role in promoting pro-inflammatory T cell signaling.

### *TYK2^D807V^* mice did not exhibit signs of autoimmune or hematological malignancies

We wanted to test if *TYK2* mutations are sufficient to alter lymphocyte function in response to cancer in genetically engineered mouse models. The amino acid sequence surrounding the D810V site in TYK2 is highly conserved between human and mouse (Extended Data Fig. 3a). Because human *TYK2^D810V^* corresponds to mouse *TYK2^D807V^*, we used CRISPR/Cas9 to generate a mouse model with *TYK2^D807V^* point mutation in all cells (Extended Data Fig. 3b-d). Germline gain-of-function mutations in JAK1-3 are commonly associated with autoimmune or hematopoietic disorders ^33–40^. Even though germline gain-of-function mutations in *TYK2* have only been reported in a few patients, we wanted to carefully survey disease development in *Tyk2^D807V^* mice.

While heterozygous (het) and homozygous (homo) mutant mice were viable and grossly indistinguishable from *TYK2^WT^* littermates (**Fig. 3a**), we evaluated whether *Tyk2^D807V^* might mediate autoimmune disease or malignancy over time. To this end, we monitored both young (2-month-old) and aged (12-month-old) cohorts. Body weights remained comparable across genotypes, with no significant weight loss or signs of cachexia observed throughout the study (**Fig. 3b**). Serum liver transaminase levels (**Fig. 3c**) and histological analysis of major organs including the liver, spleen and kidney revealed no abnormalities or inflammatory infiltration in mutant mice, supporting a lack of systemic inflammation, autoimmune manifestations, or tumorigenesis (**Fig. 3d,e** and Extended Data Fig. 4a). To assess potential hematological malignancies, we assessed the peripheral blood parameters and spleen morphology. Complete blood counts revealed no significant differences among WT, het, and homo mice (**Fig. 3f** and Extended Data Fig. 4c). Additionally, gross anatomical inspection of the spleen also showed no signs of splenomegaly or discoloration (**Fig. 3g** and Extended Data Fig. 4b). Despite JAK-STAT pathway activation in *Tyk2^D807V^* mutant mice (**Fig. 3h**), this did not drive spontaneous autoimmune pathology or oncogenic transformation. These findings suggest that *Tyk2^D807V^*, while activating, does not by itself initiate overt disease phenotypes in mice under homeostatic conditions, regardless of zygosity.

**Figure 3.**
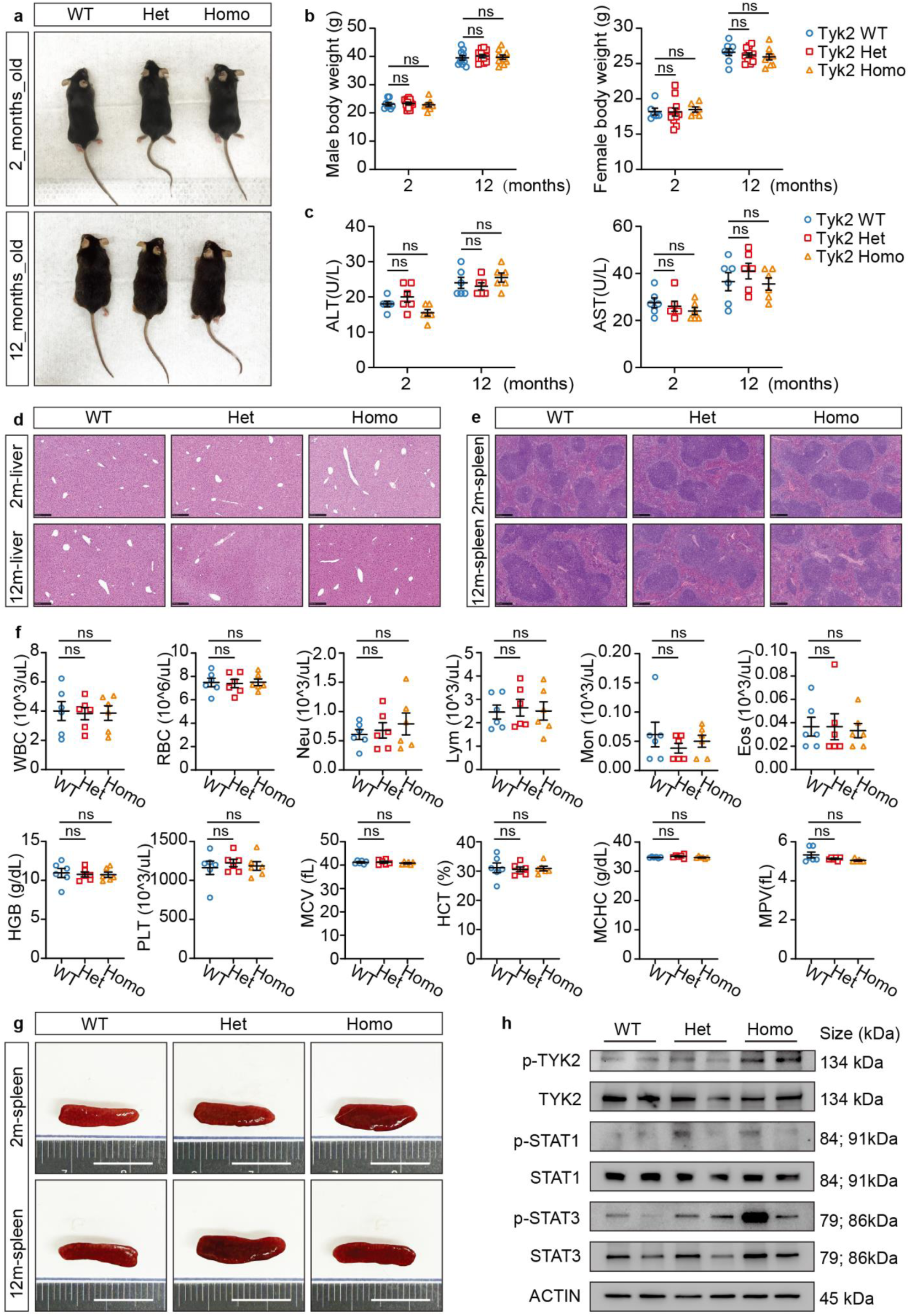
*TYK2^D807V^*mice do not exhibit signs of autoimmunity or premalignancy. **a.** Representative appearance of 2- and 12-month-old het or homo *Tyk2^D807V^* and *Tyk2^WT^* mice. **b.** Body weights of the above mice. Males: WT (*n* =8); het (*n* =12); homo (*n* =7). Females: WT (*n* =6); het (*n* =11); homo (*n* =6). **c.** Serum levels of ALT and AST (U/L) of the above mice (WT (*n* = 6); het (*n* = 6); homo (*n* = 6). mean ± s.e.m. is depicted. **d.** Representative H&E-stained liver sections from above mice (scale bars = 250 μm). **e.** Representative H&E-stained splenic sections from above mice (scale bars = 250 μm). **f.** Complete blood counts from 12-month-old mice. **g.** Representative spleens from above mice (scale bars = 10 mm). **h.** Representative western blot of liver tissues from 2-month old mice. For the data in this figure, each dot is one mouse. Error bars show mean ± s.e.m. *P* values were determined by one-way ANOVA followed by Tukey’s multiple comparisons test. \**P* < 0.05; \*\**P* < 0.01; \*\*\**P* < 0.001; \*\*\*\**P* < 0.0001; ns, not significant.

### *TYK2^D807V^* mice exhibited increased anti-tumor effects

To assess the impact of this mutation on tumor growth, we implanted syngeneic MC38 colorectal cancer cells into male *Tyk2^WT^* and *Tyk2^D807V^* mice. MC38 growth was significantly inhibited in *Tyk2^D807V^* het and homo mice vs. *Tyk2^WT^* controls (**Fig. 4a-c**). Homo mice only had a trend toward decreased tumor growth compared to het mice, indicating that one mutant allele was sufficient to cause tumor suppression. Given that patients typically harbor this somatic mutation in the het state, we used het *Tyk2^D807V^* mice for all subsequent experiments. This mutation exerted anti-tumor effects in both male and female mice (**Fig. 4d**). MC38 tumor dissociation and flow cytometry analysis revealed an increase in tumor-infiltrating CD4⁺ and CD8⁺ T cells in *Tyk2^D807V^* heterozygous mice compared to WT controls, accompanied by a reduction in myeloid subsets such as neutrophils and monocytes, and an increase in dendritic cells and NK cells (**Fig. 4e** and Extended Data Fig. 5a). Specifically, proportions of CD4⁺ effector memory T cells, CD8⁺ effector and resident T cells, and CD8⁺ effector memory T cells were elevated (**Fig. 4f**), reflecting a reshaped tumor immune microenvironment that favored adaptive immunity. A single mutant allele of *Tyk2^D807V^* mutation was sufficient to enhance T cell infiltration and suppress tumor growth in vivo.

**Figure 4.**
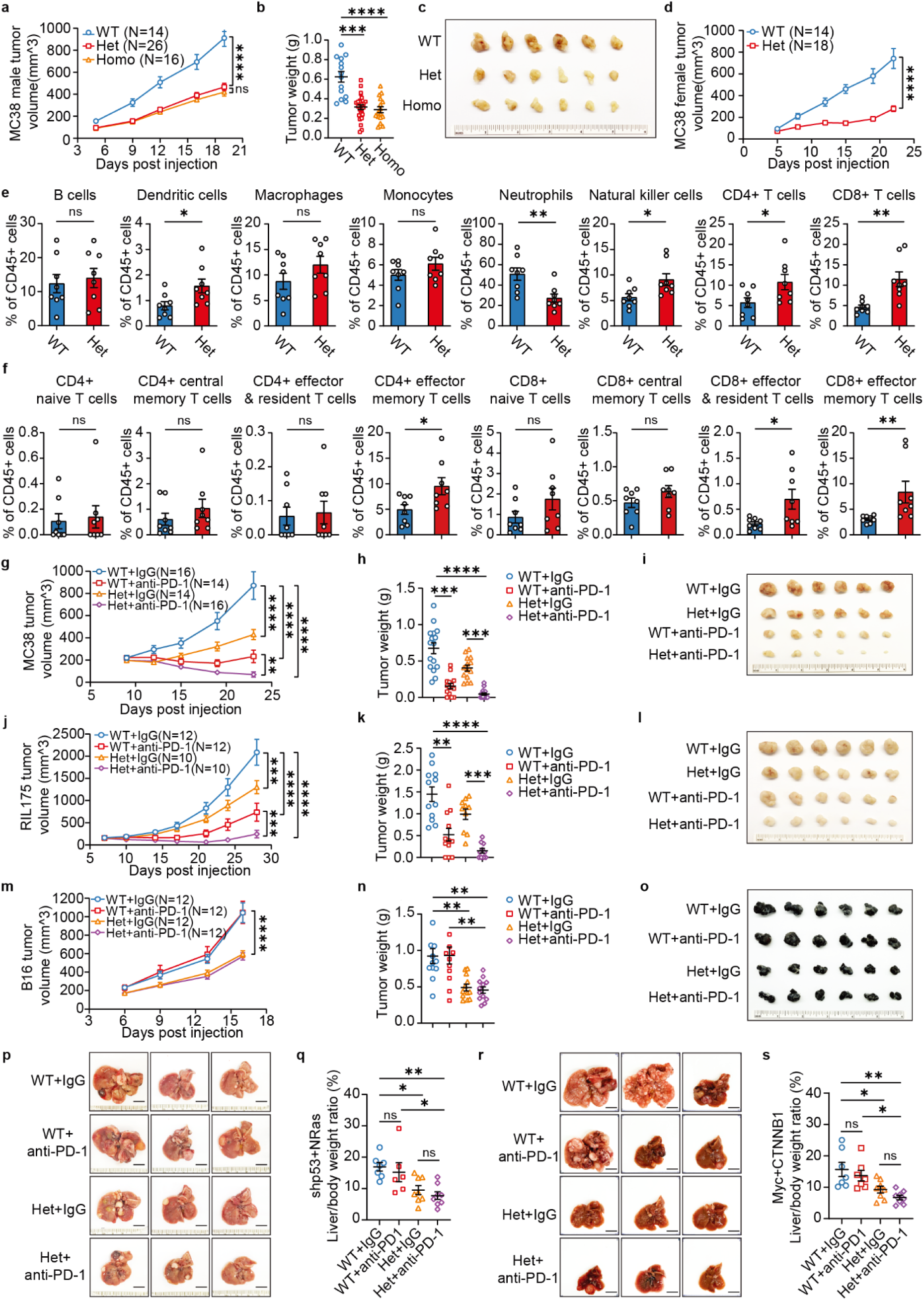
*TYK2^D807V^*mice show enhanced anti-tumor effects. **a.** Syngeneic MC38 growth in male *Tyk2^WT^*, het, or homo *Tyk2^D807V^* mice. The tumor number is shown as the “n” in the figure and two tumors per mouse were implanted. *P* values were determined by two-way ANOVA. **b.** Tumor weights from panel **a**. *P* values were determined by one-way ANOVA followed by Tukey’s multiple comparisons test. **c.** Representative tumors from each group in panel **a** were ranked by size in descending order and the top six are shown. **d.** Syngeneic MC38 growth in female het *Tyk2^D807V^* mice and *Tyk2^WT^* littermates. The tumor number is shown as the “n” in the figure and two tumors per mouse were implanted. *P* values were determined by two-way ANOVA. **e.** Quantification of tumor-infiltrating CD45⁺ cells analyzed by flow cytometry. Each data point indicates an individual tumor (WT (*n* = 8); Het (*n* = 8)). *P* values were determined using two-tailed unpaired *t*-tests. **f.** Quantification of CD4⁺ and CD8⁺ T cell subsets from tumors analyzed by flow cytometry. Each data point indicates an individual tumor (WT (*n* = 8); Het (*n* = 8)). *P* values were determined using two-tailed unpaired *t*-tests. **g.** Syngeneic MC38 growth +/- anti-PD-1 antibodies. The tumor number is shown as the “n” in the figure and two tumors per mouse were implanted. *P* values were determined by two-way ANOVA. **h.** Tumor weight collected from panel **g**. *P* values were determined by one-way ANOVA followed by Tukey’s multiple comparisons test. **i.** Representative tumors from each group in panel **g** were ranked by size in descending order and the top six are shown. **j.** Syngeneic RIL175 growth, +/- anti-PD-1 antibodies. The tumor number is shown as the “n” in the figure and two tumors per mouse were implanted. *P* values were determined by two-way ANOVA. **k.** Tumor weight collected from panel **j**. *P* values were determined by one-way ANOVA followed by Tukey’s multiple comparisons test. **l.** Representative tumors from each group in panel **j** were ranked by size in descending order and the top six are shown. **m.** Syngeneic B16 growth, +/- anti-PD-1 antibodies. The tumor number is shown as the “n” in the figure and two tumors per mouse were implanted. *P* values were determined by two-way ANOVA. **n.** Tumor weight collected from panel **m**. *P* values were determined by one-way ANOVA followed by Tukey’s multiple comparisons test. **o.** Representative tumors from each group in panel **m** were ranked by size in descending order and the top six are shown. **p.** Representative livers of *Tyk2^D807V^* het and WT littermates with shp53/*NRAS^G12V^* tumors, +/- anti-PD-1 antibodies (scale bar = 10 mm). **q.** Liver-to-body weight ratios from panel **p**. Each data point is one liver and *P* values were determined by one-way ANOVA followed by Tukey’s multiple comparisons test. **r.** Representative livers of *Tyk2^D807V^* het and WT littermates with *CTNNB1*/*MYC* tumors, +/- anti-PD-1 antibodies (scale bar = 10 mm). **s.** Liver-to-body weight ratios from panel **r**. Each data point is one liver and *P* values were determined by one-way ANOVA followed by Tukey’s multiple comparisons test. For all data in this figure, error bars show mean ± s.e.m and statistical tests are stated above. \**P* < 0.05; \*\**P* < 0.01; \*\*\**P* < 0.001; \*\*\*\**P* < 0.0001; ns, not significant.

ICIs unleash antitumor immune responses; however, only a subset of patients benefit ^41^. Given the moderate sensitivity of MC38 tumors to ICIs, we treated tumor-bearing mice with either anti-IgG control or anti-PD-1 antibodies to assess therapeutically synergistic effects of *Tyk2^D807V^* mutation. The combination led to greater inhibition compared to either intervention alone (**Fig. 4g-i**). To determine if this effect extended to other cancer types with different levels of immunogenicity, we tested the highly immunogenic RIL175 hepatocellular carcinoma (HCC) and a poorly immunogenic B16 melanoma in syngeneic settings, using *Tyk2^WT^* and *Tyk2^D807V^* mice treated with anti-IgG or anti-PD-1 antibodies. In the syngeneic RIL175 model, the combination therapy resulted in greater cancer suppression, indicating the enhancement of ICI efficacy by *Tyk2^D807V^* (**Fig. 4j-l**). In contrast, the B16 melanoma model, which is ordinarily resistant to ICIs, showed only modest tumor inhibition with anti-PD-1 treatment alone. However, *Tyk2^D807V^* mutant mice exhibited the ability to suppress B16 growth in the absence of PD-1 blockade (**Fig. 4m-o**). This suggested that the anti-tumor effect is likely independent of PD-1, but may be mediated through enhanced JAK-STAT signaling and increased IFN-γ secretion.

Syngeneic or xenograft models often fail to recapitulate the complexity of the original tumor microenvironment ^42,43^, so we examined the impact of *Tyk2^D807V^* in autochthonous models of HCC, a cancer type that responds modestly to anti-PD-1 treatments. Since *TP53* and *RAS* are among the most frequently altered pathways in HCC ^44^, we used hydrodynamic transfection (HDT) to deliver transposons carrying *NRAS^G12V^* and an shRNA against *Tp53*. This results in robust multifocal HCC development within 4-8 weeks ^45^. An important aspect of this model is that the *Tyk2^D807V^* mutation also existed within the cancer cells, and allowed us to test cell autonomous effects of the mutation. While anti-PD-1 treated *Tyk2^WT^* mice did not show significant tumor inhibition, the *Tyk2^D807V^* mutation significantly impaired tumor growth. In combination with anti-PD-1 treatment, the *Tyk2^D807V^* mutation contributed the majority of the inhibitory effect (**Fig. 4p,q**). To assess whether the phenotype is independent of the *NRAS^G12V^/TP53* oncogenic background, we utilized an alternative autochthonous HCC model based on co-expression of mutant *CTNNB1* and *MYC*, commonly mutated oncogenes in HCC ^44^. Similar to the *NRAS^G12V^*/*shp53* model, anti-PD-1 treatment had very limited efficacy, whereas *Tyk2^D807V^* significantly inhibited tumor growth. When combined with anti-PD-1, the *Tyk2^D807V^* mutation again contributed the majority of the inhibitory effect (**Fig. 4r,s**). Collectively, these findings showed that ICI efficacy can be augmented by *Tyk2^D807V^*. Remarkably in some cancer models, *Tyk2^D807V^* exerted a much more potent anti-tumor effect than anti-PD-1 antibodies.

### *Tyk2^D807V^* reduces tumor burden due to effects within CD8**⁺** T cells

In order to identify the immune cell types responsible for the anti-cancer effects exhibited by *Tyk2^D807V^* mice, we first performed immune cell depletion experiments using antibodies targeting T cells (anti-CD3ε), NK cells (anti-NK1.1), macrophages (anti-F4/80), and neutrophils (anti-Ly6G). Among these, depletion of NK cells, macrophages, or neutrophils failed to rescue syngeneic tumor growth (**Fig. 5a–f** and Extended Data Fig. 6a), suggesting that these innate myeloid immune compartments, as well as NK cells, were dispensable for the anti-tumor response of *Tyk2^D807V^*. In contrast, CD3⁺ T cell depletion abolished the anti-tumor effects of *Tyk2^D807V^* mice, implicating T cells as the primary mediators of the *Tyk2^D807V^*-dependent anti-tumor response (**Fig. 5g,h** and Extended Data Fig. 6a). To further delineate the contribution of T cell subsets, we selectively depleted CD8⁺ (anti-CD8β) or CD4⁺ (anti-CD4) T cells. The anti-tumor effect of the *Tyk2^D807V^* mutation was dependent on CD8⁺ T cells, whereas CD4⁺ T cell depletion had little impact (**Fig. 5i–l** and Extended Data Fig. 6b).

**Figure 5.**
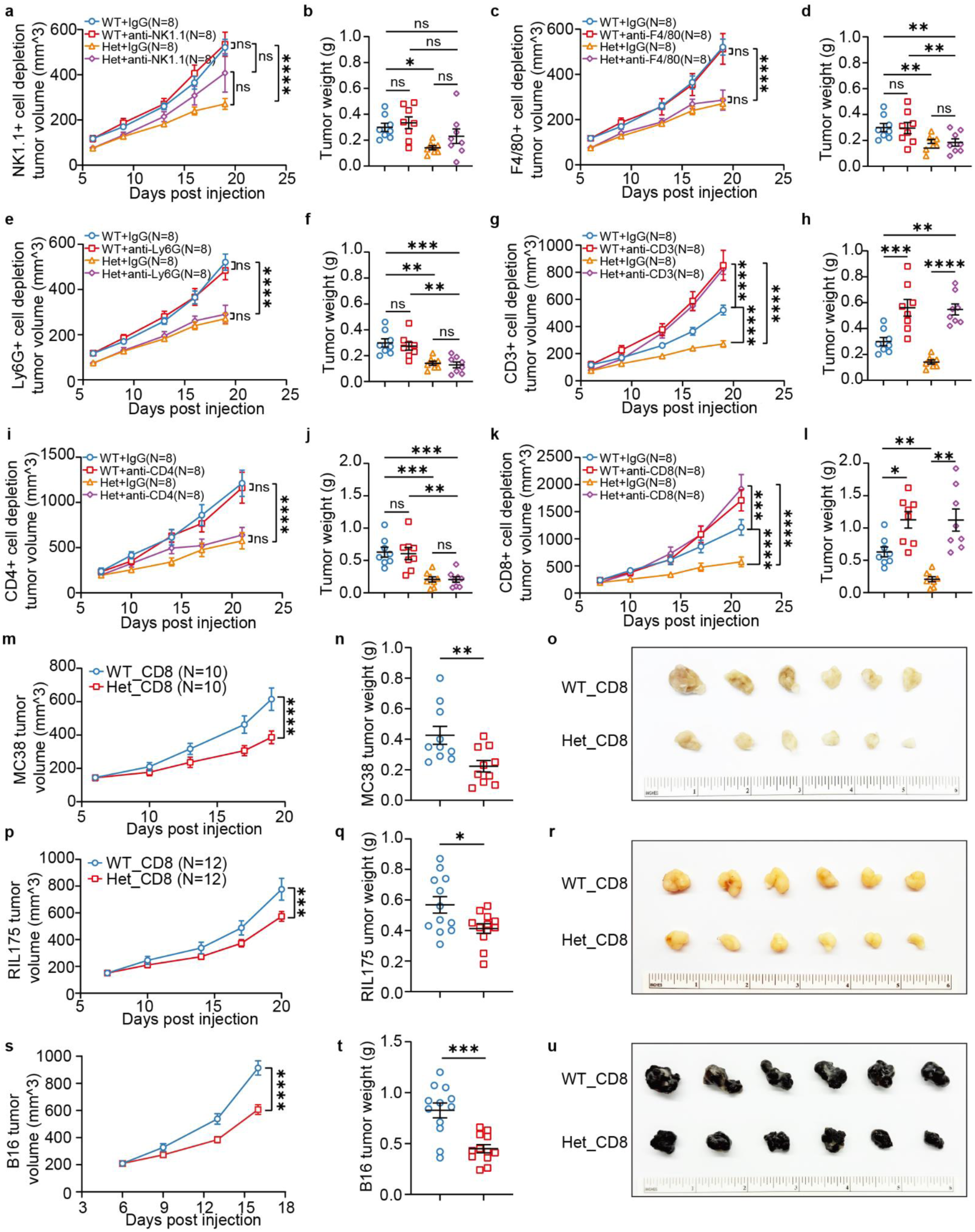
*Tyk2^D807V^*reduces tumor burden in vivo through CD8⁺ **T cells. a.** Syngeneic MC38 growth in *Tyk2^D807V^* het or WT littermates, with anti-NK1.1 or anti-IgG antibody treatments. **b.** Tumor weights from panel **a**. **c.** Syngeneic MC38 growth in *Tyk2^D807V^* het or WT littermates, with anti-F4/80 or anti-IgG antibody treatments. **d.** Tumor weights from panel **c**. **e.** Syngeneic MC38 growth in *Tyk2^D807V^* het or WT littermates, with anti-Ly6G or anti-IgG antibody treatments. **f.** Tumor weights from panel **e**. **g.** Syngeneic MC38 growth in *Tyk2^D807V^* het or WT littermates, with anti-CD3 or anti-IgG antibody treatments. **h.** Tumor weights from panel **g**. **i.** Syngeneic MC38 growth in *Tyk2^D807V^* het or WT littermates, with anti-CD4 or anti-IgG antibody treatments. **j.** Tumor weights from panel **i**. **k.** Syngeneic MC38 growth in *Tyk2^D807V^* het or WT littermates, with anti-CD8 or anti-IgG antibody treatments. **l.** Tumor weights from panel **k**. **m.** Syngeneic MC38 growth in mice transplanted with CD8⁺ T cells from *Tyk2^D807V^* het or WT littermates. **n.** Tumor weights from panel **m**. **o.** Representative tumors from panel **m**. **p.** Syngeneic RIL175 growth in mice transplanted with CD8⁺ T cells from *Tyk2^D807V^* het or WT littermates. **q.** Tumor weights from panel **p**. **r.** Representative tumors from panel **p**. **s.** Syngeneic B16 growth in mice transplanted with CD8⁺ T cells from *Tyk2^D807V^* het or WT littermates. **t.** Tumor weight collected from panel **s**. **u.** Representative tumors from panel **s**. All data in this figure are presented as mean ± s.e.m. Tumor cells were subcutaneously injected into both flanks of each mouse, so each mouse bears two tumors. Each data point represents an individual tumor (N). For syngeneic models growth curves (panels **a,c,e,g,i,k,m,p,s**), *P* values were determined by two-way ANOVA. For tumor weight panels, *P* values were determined by one-way ANOVA followed by Tukey’s multiple comparisons test (panels **b,d,f,h,j,i**), or two-tailed unpaired *t*-tests (panels **n,q,t**). \**P* < 0.05; \*\**P* < 0.01; \*\*\**P* < 0.001; \*\*\*\**P* < 0.0001; ns, not significant. Of note, the data in panels **a**-**h** share the same WT+anti-IgG and het+anti-IgG control groups. And the data in panels **i-l** share the same WT+anti-IgG and het+anti-IgG control groups. For all gross images, tumors from each group were ranked by size in descending order, and the top six are shown.

To determine if *Tyk2^D807V^* CD8⁺ T cells were sufficient to suppress established tumors, we performed adoptive T cell transfer experiments. CD8⁺ T cells were isolated from either *Tyk2^WT^* or *Tyk2^D807V^* het mice and transferred into immunocompetent WT recipients bearing established MC38 syngeneic tumors (Extended Data Fig. 6c). Notably, recipients of *Tyk2^D807V^* CD8⁺ T cells exhibited significantly delayed tumor growth and reduced tumor burden compared to those receiving *Tyk2^WT^* CD8⁺ T cells, indicating that *Tyk2^D807V^* intrinsically enhances CD8⁺ T cell effector function (**Fig. 5m–o**). To test generalizability, we evaluated additional cancer models. In both the RIL175 (**Fig. 5p–r**) and B16 (**Fig. 5s–u**) models, transfer of *Tyk2^D807V^* mutant CD8⁺ T cells consistently inhibited tumor growth. These results revealed that the *Tyk2^D807V^* mutation conferred CD8⁺ T cells with a broadly effective anti-tumor program, capable of overcoming the immune resistance of poorly immunogenic tumors, and suggested a potential strategy for engineering hyperactive cytotoxic T cells for adoptive immunotherapy.

### *Tyk2^D807V^* also impaired cancer growth in a cell-autonomous manner

One potential negative aspect of mutating *Tyk2* in all cells in the body is that TYK2 activation and increased JAK-STAT signalling could have tumor intrinsic growth effects. In the above HDT experiments, increased T cell activities could have obscured a cell autonomous pro-tumor effect of TYK2 activation. Previously, hyperactive TYK2, JAK3, and STAT3 signaling within cancer cells have been shown to promote cancer growth ^38–40,46^, an observation that would have to be considered if *Tyk2* activation is used to treat cancer. To test if *Tyk2^D807V^* activity within tumors promoted HCC growth independently of *Tyk2^D807V^* activity within immune cells, we isolated *NRAS^G12V^*/*shp53* HCC cells from *Tyk2^WT^* and *Tyk2^D807V^* het mice. In vitro, these HCC cells did not exhibit differences in growth (**Fig. 6a**). When implanted into WT C57BL/6 mice, *Tyk2^D807V^* HCCs surprisingly showed impaired tumor growth and reduced tumor burden compared to *Tyk2^WT^* HCCs ( **Fig. 6b-d**).

**Figure 6.**
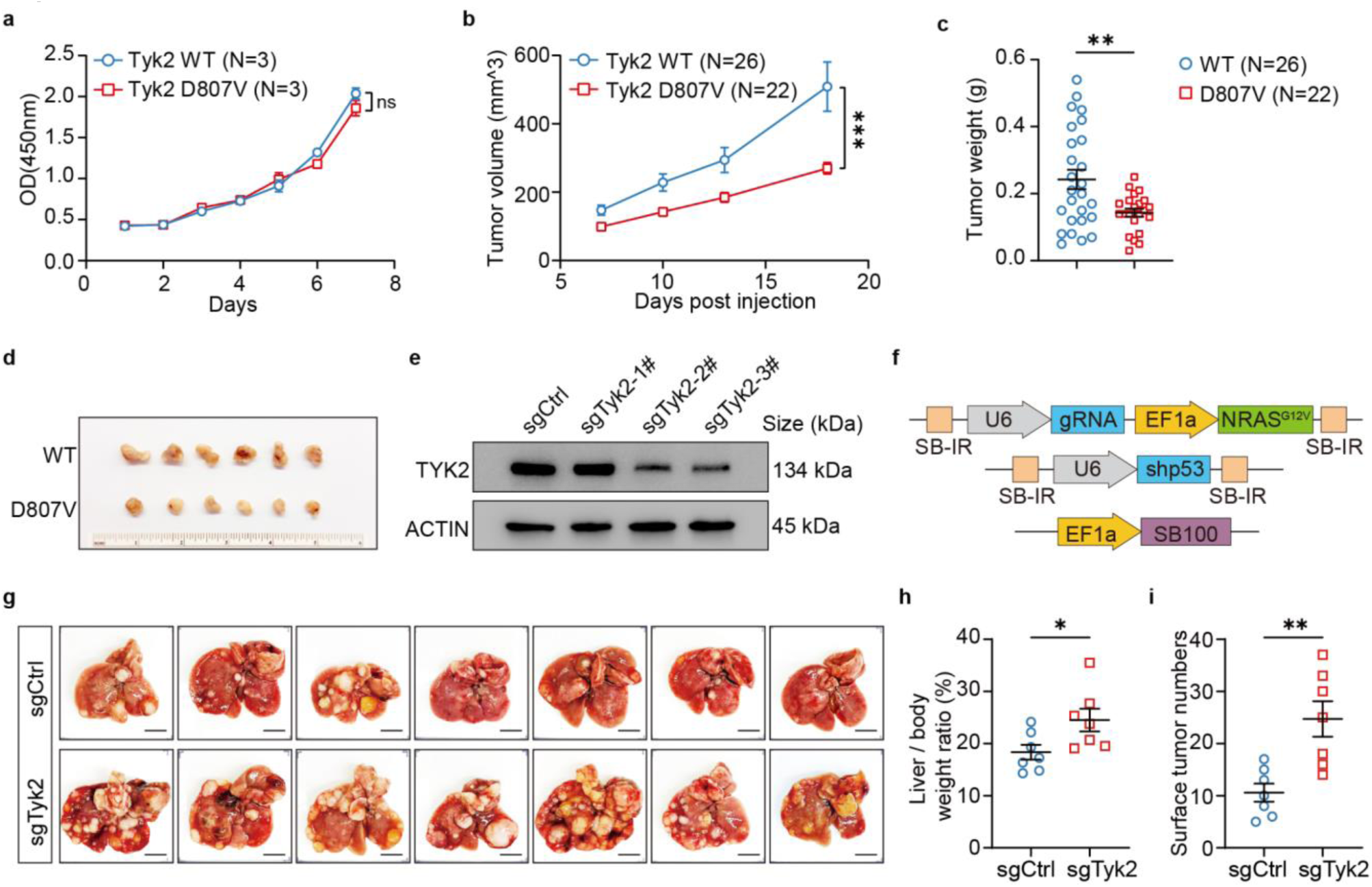
*Tyk2^D807V^*also impairs cancer growth in a cell-autonomous manner. **a.** Cell viability assay (CCK-8) performed on *NRAS^G12V^*/*shp53* HCC cells isolated from WT and *Tyk2^D807V^* het mice grown in culture (*n* = 3,3). *P* values were determined by two-way ANOVA. **a. b.** *NRAS^G12V^*/*shp53* HCC cells isolated from WT and *Tyk2^D807V^* het mice were implanted into WT C57/B6 mice (*n* = 26 *Tyk2^WT^* and 22 *Tyk2^D807V^* het tumors). **b.** Tumor weights from panel **b**. *P* values were determined using two-tailed unpaired *t*-test. **c.** Representative tumors from panel **b**. Tumors were ranked by size in descending order, and the top six are shown. **d.** Western blot of H2.35 cells overexpressing sg*TYK2* knockout vectors. **e.** Transposon constructs containing sgRNA-*NRAS^G12V^*, *shp53*, and a plasmid containing *SB100* transposase. **f.** Representative livers of sg*Tyk2* and *sgCtrl* with shp53/*NRAS^G12V^* tumors (scale bar = 10 mm). **g.** Liver-to-body weight ratios from panel **g**. Each data point is one liver and *P* values were determined by two-tailed unpaired *t*-test. **h.** Surface tumor numbers from panel **g**. Each data point is one liver and *P* values were determined by two-tailed unpaired *t*-test. All data in this figure are presented as mean ± s.e.m. Tumor cells were subcutaneously injected into both flanks of each mouse, so each mouse bears two tumors. Each data point represents an individual tumor (N). \**P* < 0.05; \*\**P* < 0.01; \*\*\**P* < 0.001; \*\*\*\**P* < 0.0001; ns, not significant.

To test the essentiality of *Tyk2* using loss of function approaches, we generated *Tyk2* KO *NRAS^G12V^*/*shp53* HCCs using HDT (**Fig. 6e-f**). In contrast with *Tyk2^D807V^*, *Tyk2* KO tumors exhibited accelerated growth, further supporting a cell intrinsic tumor-suppressive role for TYK2 in vivo (**Fig. 6g–i**). These data provide compelling evidence that TYK2 activation in TILs enhances anti-tumor immunity, while activation in tumor cells inhibits cancer progression–thus providing a dual rationale for TYK2 activation across tissues.

### Pharmacological inhibition of TYK2 promotes tumor growth

The observation that *Tyk2^D807V^* can increase anti-cancer immunity non-cell autonomously and decrease cancer growth cell autonomously suggests that under non-mutant conditions, WT TYK2 activity could also restrain tumor growth. Deucravacitinib selectively targets the TYK2 pseudokinase domain and is used clinically to block JAK-STAT in autoimmune diseases ^27–29^. Given that the *TYK2^D807V^* mutation also occurs in the pseudokinase domain and activates the JAK-STAT pathway, we hypothesized that treatment with deucravacitinib could impair anti-cancer immunity. In vitro, deucravacitinib significantly impaired T cell–mediated killing of A549 tumor cells in a co-culture assay (**Fig. 7a,b**). Consistently, IFN-γ levels in co-culture supernatants were decreased (**Fig. 7c**), indicating suppressed T cell effector function following deucravacitinib treatment.

**Figure 7.**
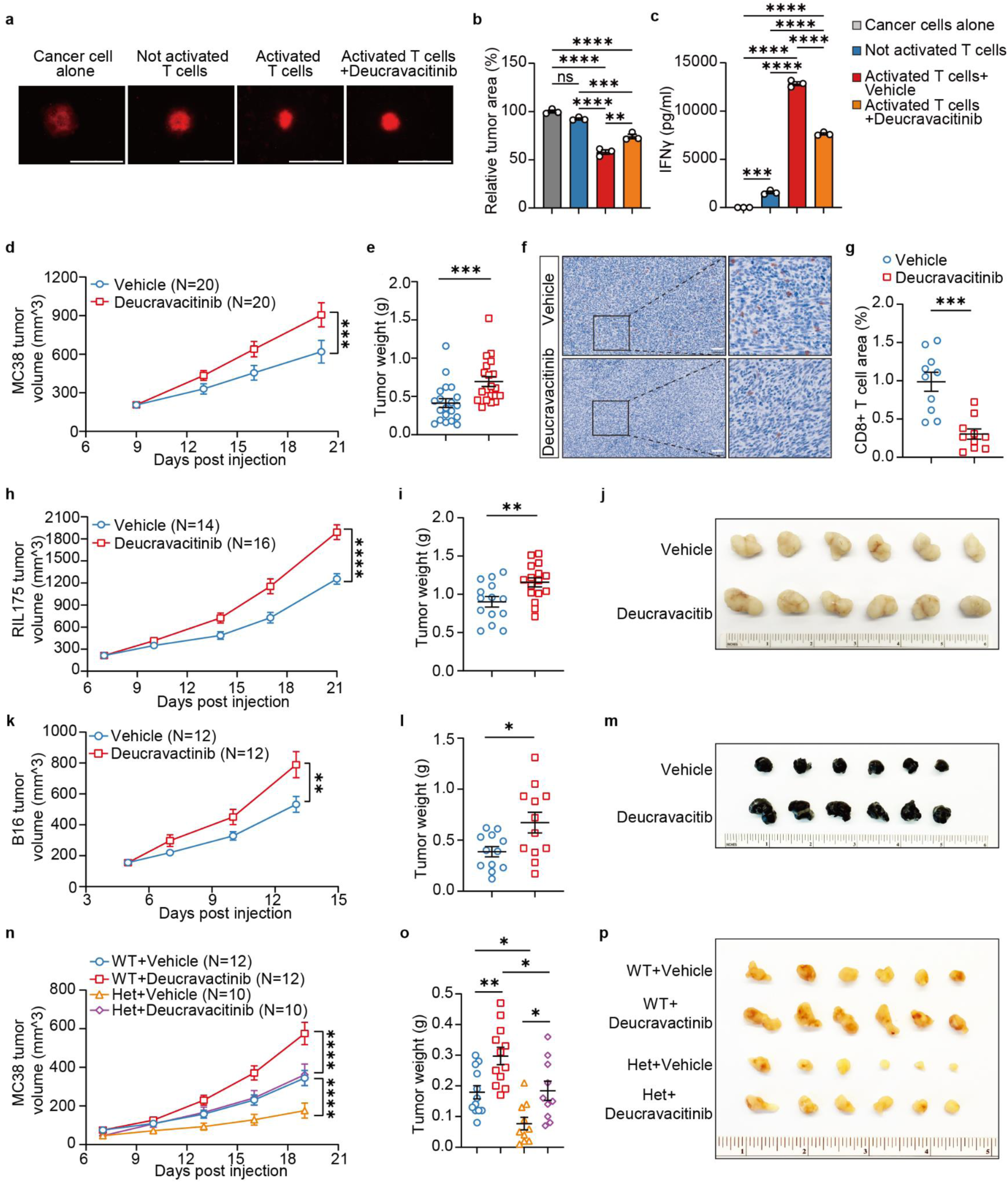
Pharmacological inhibition of TYK2 promotes tumor growth. **a.** Representative images of A549 tumor co-cultured with primary T cells after treatment with Deucravacitinib or vehicle, scale bars = 1000 μm (*n=*3 replicates per group). **b.** Relative tumor area of A549 tumor co-cultured with primary T cells after treatment with Deucravacitinib or vehicle (*n*=3 replicates per group). **c.** IFNγ levels measured in A549 tumor cells co-cultured with primary T cells after treatment with Deucravacitinib or vehicle (*n*=3 replicates per group). **d.** Syngeneic MC38 growth in mice receiving deucravacitinib (*n*=16 tumors) or vehicle control (*n*=14 tumors). **e.** Tumor weight collected from panel **d**. **f.** Representative IHC images of tumor infiltrating CD8⁺ cells in MC38 implanted tumors receiving deucravacitinib or vehicle. (scale bars = 150 μm) **g.** Quantification of tumor infiltrating CD8⁺ cell area of panel **f** (*n*=10,10 for vehicle, deucravacitinib treated tumors). **h.** Syngeneic RIL175 tumor growth in mice receiving deucravacitinib (*n*=14 tumors) or vehicle control (*n*=16 tumors). **i.** Tumor weight collected from panel **h**. **j.** Representative syngeneic tumors from panel **h**. Tumors from each group were ranked by size in descending order, and the top six were selected for imaging. **k.** Syngeneic B16 tumor growth in mice receiving deucravacitinib (*n*=12 tumors) or vehicle control (*n*=12 tumors). **l.** Tumor weight collected from panel **k**. **m.** Representative syngeneic tumors from panel **k**. Tumors from each group were ranked by size in descending order, and the top six were selected for imaging. **n.** Syngeneic MC38 growth in *Tyk2^D807V^* het or WT littermates, with vehicle or deucravacitinib treatments. **o.** Tumor weights from panel **n**. **p.** Representative syngeneic tumors from panel **n**. Tumors from each group were ranked by size in descending order, and the top six were selected for imaging. All data in this figure are presented as mean ± s.e.m. Tumor cells were subcutaneously injected into both flanks of each mouse, so each mouse bears two tumors. Each data point represents an individual tumor (N). For syngeneic models growth curves (panels **d,h,k,n**), *P* values were determined by two-way ANOVA. For panels **e,g,i,l,o**, *P* values were determined by two-tailed unpaired *t*-tests. \**P* < 0.05; \*\**P* < 0.01; \*\*\**P* < 0.001; \*\*\*\**P* < 0.0001; ns, not significant.

Next, we evaluated the in vivo impact of pharmacological TYK2 inhibition across multiple tumor models. Deucravacitinib was administered to immunocompetent C57BL/6 WT mice bearing MC38, RIL175, or B16 syngeneic tumors. In all three models, deucravacitinib significantly accelerated tumor growth compared to vehicle controls and was accompanied by a marked reduction in intratumoral CD8⁺ T cell infiltration (**Fig. 7d–m**). These findings indicated that TYK2 inhibition broadly impairs anti-tumor immunity across diverse tumor types. To determine if deucravacitinib could antagonize the tumor-suppressive effects of *Tyk2^D807V^*, we treated MC38 tumor-bearing *Tyk2^D807V^* mice with vehicle or deucravacitinib. This attenuated the anti-cancer effects of *Tyk2^D807V^*, but did not fully abrogate the phenotype. Instead, tumor growth increased to *Tyk2^WT^* levels, suggesting that the dose or mechanism of deucravacitinib is insufficient to fully counteract *Tyk2^D807V^*’s anti-tumor effects (**Fig. 7n–p**). While TYK2 blockade may offer therapeutic benefits in autoimmune diseases, such interventions could potentially exacerbate tumor growth or undermine anti-cancer immune surveillance in certain settings.

## Discussion

We identified somatic mutations in TILs from cancer patients and demonstrated their ability to enhance T cell-mediated immune responses. A critical limitation of somatic genomics up to this point is that sequencing studies have focused on known cancer associated genes that occur at a high enough VAF to be detected with standard sequencing approaches. As a result, many genes and mutations not typically linked to cancer, especially those with low VAFs, have been overlooked. An important aspect of our study is the application of NanoSeq, a highly sensitive genome-wide duplex sequencing method that enables accurate detection of rare mutations in all coding genes, including those occurring at extremely low VAFs in small populations of non-malignant cells.

Among the mutations identified, those affecting *TYK2* were of particular interest. These mutations not only enhance T cell effector function but may also hold potential as therapeutic targets in cancer treatment. A key observation here is that these mutations do not lead to autoimmunity at baseline but enhance T cell activation in the context of cancer. Unlike oncogenic mutations commonly found in T cell neoplasms, which can contribute to transformation, the *TYK2* mutations identified in non-neoplastic lymphocytes appear to support T cell function without inducing pathogenic outcomes. Mimicking mutations in non-neoplastic T cells presents an opportunity to enhance immune responses against tumors without increasing the risk of malignant transformation. Our findings resonate with recent work describing somatic loss-of-function mutations in negative regulators such as *TNFAIP3* in TILs, which support a model where removing inhibitory constraints enhances antitumor immunity ^16^. However, *TNFAIP3* is a well known tumor suppressor in various lymphomas, which might increase the risk associated with therapeutic manipulation ^47^.

Importantly, our findings have implications for cancer immunotherapy. The *Tyk2^D810V^* mutation within T cells may synergize with ICIs to potentiate anti-tumor responses. Notably, this mutation may also provide therapeutic benefit in tumors resistant to ICIs, suggesting that engineering T cells to express *Tyk2^D810V^* could deliver robust cytotoxicity independent of ICI therapy. This unexpected efficacy likely arises because the *Tyk2^D810V^* mutation is adaptively selected within the tumor microenvironment of solid tumors, including poorly immunogenic melanomas, to enable T cells to overcome local immunosuppression. Accordingly, immunotherapies harnessing such naturally occurring somatic mutations in TILs could improve clinical outcomes in patients with solid tumors typically refractory to checkpoint blockade.

In summary, our study highlights the importance of understanding the functional consequences of somatic mutations outside of cancer, but within the tumor microenvironment. While appropriate attention has been given to the mutations in cancer cells, our work emphasizes the value of exploring naturally occurring mutations within non-neoplastic immune cells. This approach provides a way of discovering unexpected immunotherapy targets. Future research will need to address several critical questions, including the optimal means of replicating the effects of these mutations in clinical settings and understanding potential interactions with other immune cell types. Our findings lay a foundation for the development of novel immunotherapies that exploit naturally occurring somatic mutations to enhance anti-cancer immunity.

## Author contributions

Z.L. and H.Z. designed and performed experiments, and wrote the manuscript. M.H.H assisted with in vivo cancer models. J.E. conceived and analyzed TYK2 experiments. A.A. and T.L. performed in vitro experiments. G.K., A.R.J.L., P.A.N., and H.E.M. generated and analyzed NanoSeq mutation data from human TILs. A.A.K. performed and analyzed scRNA-seq on human lymphocytes. R.W. supervised human TYK2 experiments. Q.Z. performed primary tumor isolation and in vitro assays. N.J. performed murine blood testing. X.W. and G.M. performed FACS analysis. X.R. assisted with cell culture. T.S. and S.K. generated the mouse models. X.F. performed histologic analysis and IHC. D. H. designed experiments and edited the manuscript. S.H., S.B., I.M., P.C. supervised genomic analysis and edited the manuscript.

## Declaration of Conflicts

H.Z. I.M., and P.C. are co-founders of Quotient Therapeutics. J.E., A.A., T.L., G.K., H.E.M., A.A.K., and R.W. were employees of Quotient Therapeutics at the time of the study. H.Z. is an advisor for Newlimit, Alnylam Pharmaceuticals, and Chroma Medicines. A provisional patent related to this work (Provisional Patent Application No. 63/886,777: ENGINEERED T CELL POPULATIONS AND RELATED METHODS) was submitted on September 23, 2025. The inventors were: Hao Zhu, Zhijie Li, Inigo Martincorena, Peter Campbell, Andrew Robert James Lawson, John Evans, Rui Wang, Xiaoyi Jin, Alisa Arutyunova, Tao Liu, Gerda Kildisiute, Matthew Young, Zeynep Kalender-Atak, Scott Hayton, Simon Brunner.

## Materials and methods

### DNA extraction of human TILs

TILs isolated from human tumours of 46 donors (including 18 lung cancer, 16 colorectal cancer, 7 kidney cancer, and 5 melanoma) were sourced from a commercial provider (Discovery Life Sciences). Lymphocyte subsets from 20 patients sequenced by the Sanger Institute included CD4 T cells, CD8 T cells, NK cells, and B cells, with a total of 75 samples, while lymphocyte subsets from 26 patients sequenced by Quotient Therapeutics included CD4 and CD8 T cells, with a total of 78 samples. Samples underwent DNA extraction using the QIAGEN QIAamp DNA Micro Kit (catalogue number 56304). DNA extraction was performed according to the manufacturer’s protocol with the following modifications: all centrifuge steps were completed at 20,000 × g, buffer AE was replaced with buffer EB, the final elution volume was 130µL.

### NanoSeq

Sequencing data from Sanger was processed using the NanoSeq pipeline (https://github.com/cancerit/NanoSeq). Resulting somatic variants were used as an input to dnds algorithm to detect positive selection at nucleotide resolution using the site model implemented in the sitednds function of the dndscv R package (https://github.com/im3sanger/dndscv). This model tests each genomic site individually for a statistically significant excess of mutations, relative to a neutral expectation. The expected mutation rates at each site were derived from the context-dependent substitution spectrum. A global background model was fitted across all synonymous sites using a negative binomial distribution with a shared overdispersion parameter and a single intercept term. Sites exhibiting a statistically significant excess of mutations (q < 0.1, FDR-corrected) were classified as being under positive selection. In order to include NanoSeq data from Quotient Therapeutics in our analysis, sequencing data from both cohorts were re-processed using proprietary versions of the NanoSeq pipeline. Resulting variants were used as an input to a proprietary implementation of dnds algorithm. Sites exhibiting a statistically significant excess of mutations (q < 0.1, FDR-corrected) were classified as being under positive selection.

### T cell activation and CRISPR perturbation

CD3⁺ T cells were isolated from peripheral blood mononuclear cells (PBMCs) of four healthy adult donors. Isolated T cells were stimulated overnight with anti-CD3/CD28 Dynabeads. Cells were then transduced with a pooled lentiviral sgRNA library comprising 184 total guides, including the following four targeting *TYK2*: 5’-GTAGTGCTGCGCGTGCAGAT; 5’-GTGCGTCAGCAGCTCATACA; 5’-GTCAAAGCAGATCTCCAGGA; 5’-GTCATGGAGTACGTGCCCCT. Cultures were expanded for five days then re-stimulated with fresh anti-CD3/CD28 beads overnight before single-cell capture.

### Single-cell encapsulation and sequencing

Approximately 105,000 viable cells were loaded per channel on a Chromium X (10x Genomics 3′ chemistry, six channels in total). Gene-expression and guide-capture libraries were prepared according to the manufacturer’s protocol and pooled equimolarly. Sequencing was performed on an Illumina NovaSeq X Plus.

### T cell cytokine production

To assess activation-induced cytokine production, 1×10⁵ T cells (Zenbio) were stimulated with a CD3/CD28 antibody cocktail (Stemcell, Cat. #10971) following the manufacturer’s instructions. Where indicated, wells were additionally treated with either 20 ng/mL IL-12, 200nM deucravacitinib (MedChemExpress HY-117287), or vehicle. After 24 hours, culture supernatants were collected and stored at –80 °C. Prior to analysis, samples were thawed on ice and diluted 1:4. IFN-γ levels were then quantified using the U-Plex MSD kit (Cat. #K15067M-1) according to the manufacturer’s protocol.

### Primary T cell transduction

Activated primary T cells were transduced with different lentiviral vectors at a multiplicity of infection of 50 through spinoculation at 2000× g for 60 minutes at 32 °C. The percentage of transduced cells was determined by flow cytometry, and the cells were sorted on an Aria III to obtain a pure population of GFP-expressing cells.

### In vitro tumor growth assay

A549 cells (ATCC, Cat. CRM-CCL-185) were seeded in ultra-low attachment plates (Corning, CLS7007) at a density of 3,000 cells per well in 2% Matrigel (Corning 354230). A single spheroid was formed in each well, after which T cells were added at effector-to-target (E:T) ratios of 10:1. Spheroid sizes were measured at 72 hours using the automated imaging system Cytation 5. Supernatants were collected for cytokine production analysis using V-Plex MSD kit (Cat. K150449D-4).

### Mice

All mice were handled in accordance with the guidelines of the Institutional Animal Care and Use Committee at UT Southwestern. All experiments were done in an age and sex controlled fashion unless otherwise noted. C57BL/6 strain background mice were used for all experiments. The *Tyk2^D807V^* mutation mice were generated using CRISPR/Cas9.

### Cell culture

The MC38 cell line was obtained from ATCC. The RIL175 cell line was kindly provided by Dr. Yujin Hoshida from UTSW. The B16 cell line was a kind gift from Dr. Lingjie Sang from UTSW. B16 cells were cultured in RPMI-1640 with 10% FBS and penicillin-streptomycin, and other cells were cultured in DMEM with 10% FBS and penicillin-streptomycin at 37°C in a 5% CO2 incubator.

### Hydrodynamic transfection (HDT)

For shP53/*NRAS^G12V^* and Myc/CTNNB1 HDT models, 10 ug/ml transposon plasmid and 0.5 ug/ml SB100 transposase were prepared in saline. The injection volume was determined according to mouse body weight (1 ml/10g). The injection was performed over 6-8 seconds through the tail vein. After injection, mice were placed on a heating pad overnight for recovery.

### Histology and immunohistochemistry (IHC)

Tissues were fixed overnight at 4°C in 4% paraformaldehyde (PFA; Alfa Aesar #J19943K2), paraffin-sectioned, and H&E stained (UTSW Histopathology Core). For IHC, paraffin-sectioned slides were deparaffinized in xylene and rehydrated in 100%, 90%, 80%, 70%, 50%, 30% ethanol and deionized water. Antigen retrieval was performed in 10 mM Sodium Citrate buffer (pH 6.0) with 0.05% Tween 20 at a sub-boiling temperature for 20 minutes in a microwave. After cooling down, slides were immersed in 3% hydrogen peroxide in methanol to block endogenous peroxidase activity. Immunohistochemistry was then performed using VECTASTAIN Elite ABC-HRP kits (Vector Laboratories #PL-6101, #PK-2200; #PK-6104) as described in the manufacturer’s instructions. The following primary antibodies and dilutions were used for immunohistochemical staining: CD8 (CST #98941S). The slides were counterstained with hematoxylin (Vector Laboratories #NC9788954).

### Western blot analysis

Frozen liver tissues or cells were treated with RIPA lysis and extraction buffer (Life Technologies #89900) supplemented with protease and phosphatase inhibitor cocktail (Life Technologies #78440). Concentration of protein lysis was quantified with Pierce BCA Protein Assay kit (Thermo Fisher Scientific #23225). The membranes were blocked with 5%(w/v) skim milk (BD Bioscience, USA) for 1 h at room temperature, and then incubated with the following primary antibodies overnight at 4 °C: TYK2 (Proteintech #16412-1-AP, 1:1000), TYK2 (Cell Signaling Technology #35615S, 1:1000), Phospho-TYK2 (Cell Signaling Technology, 68790S, 1:1000), STAT1 (Cell Signaling Technology, 14994S, 1:1000), Phospho-STAT1 (Cell Signaling Technology, 9167S, 1:1000), STAT3 (Cell Signaling Technology, 9139T, 1:1000), Phospho-STAT3 (Cell Signaling Technology, 9145S, 1:1000) and β-actin (Cell Signaling #4970, 1:1000).

### Syngeneic cancer models

1×10^6^ MC38, 0.5×10^6^ RIL-175, or 1×10^5^ B16 cells were suspended in 50% matrigel (Corning Life Sciences) and 50% Hank’s Balanced Salt Solution (HBSS, Sigma). Cells were subcutaneously implanted into the flanks of 6-8 week old mice. Tumor volumes were measured twice per week using electronic calipers and estimated by the modified ellipsoid formula: tumor volume = (length x width^2^) / 2.

### Immune cell depletion in mice

For macrophage and monocyte depletion, anti-F4/80 (Bio X cell #BE0206) antibodies were administered at 200 µg/mouse daily for 5 days. All other antibodies including anti-CD3ε (Bio X cell #BE0001-1FAB), anti-Ly6G (Bio X cell #BP0075-1), anti-NK1.1 (Bio X cell #BE0036), anti-CD4 (Bio X cell #BP0003-1), and anti-CD8β (Bio X cell #BE0223) were administered at 200 µg/mouse twice per week.

### Liver function tests

Blood was taken using heparinized tubes from the inferior vena cava immediately after sacrificing the mouse, and then transferred into 1.5 ml tubes and centrifuged at 2000 g for 15 minutes at 4°C. The supernatant after centrifugation (plasma) was analyzed for AST and ALT (Manufacturer’s Reference Numbers 8433815 and 1655281, respectively) using a fully automated OCD Vitros 350 dry chemistry analyzer following the protocols provided by the reagent kit manufacturer (Ortho Clinical Diagnostics, Raritan, NJ) at the UT Southwestern Metabolic Phenotyping Core.

### RNA extraction and transcriptome analysis in Jurkat T cells

Total RNA from Jurkat T cells was isolated using TRIzol reagent (Invitrogen #15596018) followed by purification using the RNeasy Mini kit (QIAGEN). For transcriptome sequencing, 1 μg of total RNA was used for the construction of sequencing libraries and the transcriptome sequencing was performed using the high throughput sequencing platform of Illumina.

### Flow cytometry analysis of mouse TILs

Tumors were excised from mice and immediately placed in ice-cold phosphate-buffered saline (PBS). Under sterile conditions, tumor tissues were minced into approximately 1–3 mm^3^ fragments and enzymatically digested in RPMI medium supplemented with 100 U/ml Collagenase IV (Life Technologies #17104019) and 0.1 mg/ml DNaseI (Sigma Aldrich #10104159001) at 37°C for 30 minutes with gentle agitation. The resulting cell suspensions were filtered through a 70 µm cell strainer to obtain single-cell suspensions. Red blood cells were lysed using ACK lysis buffer (ThermoFisher Scientific) for 5 minutes at room temperature, followed by washing with PBS. For flow cytometry, cells were resuspended in FACS buffer (PBS containing 0.5% fetal bovine serum) and stained with following fluorochrome-conjugated antibodies: CD45 (Biolegend #100310), B220 (Biolegend # 103232), NK1.1 (Biolegend # 108708), CD11b (Invitrogen # 17-0112-83), CD11c (Biolegend # 117306), Ly6C (BD Biosciences # 562727), Ly6G (BD Biosciences # 560601), CD3 (Biolegend # 100204), CD4 (Invitrogen # 25-0041-82), CD8a (Tonbo # 20-0081-U100), CD44 (Biolegend # 103026), CD62L (Biolegend # 104408), TER-119 (BD Biosciences # 563995).

### Statistical analysis

Mice were randomly allocated to experiments, and samples were processed in an arbitrary order; formal randomization techniques were not applied. Data were first assessed for normality using the Shapiro–Wilk test (for sample sizes 3 ≤ n < 20) or the D’Agostino–Pearson omnibus test (for n ≥ 20). To test whether variability significantly differed among groups, we performed *F*-tests (for experiments with two groups) or Brown-Forsythe test (for experiments with more than two groups). When data deviated significantly from normality or variability significantly differed among groups, log₂ transformation was applied, followed by normality and variability test again. If the transformed data no longer significantly deviated from normality and equal variability, parametric tests were performed on transformed data; otherwise, non-parametric tests were used on original data. For two-group comparisons, if both assumptions of normality and equal variances were met, unpaired two-tailed Student’s t-tests were used. If variances significantly differed, Welch’s t-test was applied. When data significantly deviated from normality and could not be normalized by log₂ transformation, the non-parametric Mann–Whitney U test was used. All statistical tests were two-sided. For comparisons involving more than two groups, if both assumptions of normality and equal variances were met, one-way ANOVA followed by Tukey’s multiple comparisons test was performed. If variances significantly differed, Welch’s ANOVA followed by Dunnett’s T3 multiple comparisons test was used. When data significantly deviated from normality and could not be normalized by log₂ transformation, the non-parametric Kruskal–Wallis test followed by Dunn’s multiple comparisons test was performed. For longitudinal measurements of syngeneic tumor volume across multiple timepoints and treatment groups, repeated-measures two-way ANOVA was performed to evaluate the effects of time, treatment, and their interaction. A mixed-effects model with the Geisser–Greenhouse correction was applied to account for within-subject correlations and potential violations of the sphericity assumption. Statistical analyses were conducted using GraphPad Prism (version 10.1.2), and a p-value < 0.05 was considered statistically significant. Statistical significance in figures is summarized as follows: \**P* < 0.05; \*\**P* < 0.01; \*\*\**P* < 0.001; \*\*\*\**P* < 0.0001; ns, not significant. Results are expressed as mean ± s.e.m.

---

**Extended data figure 1.**
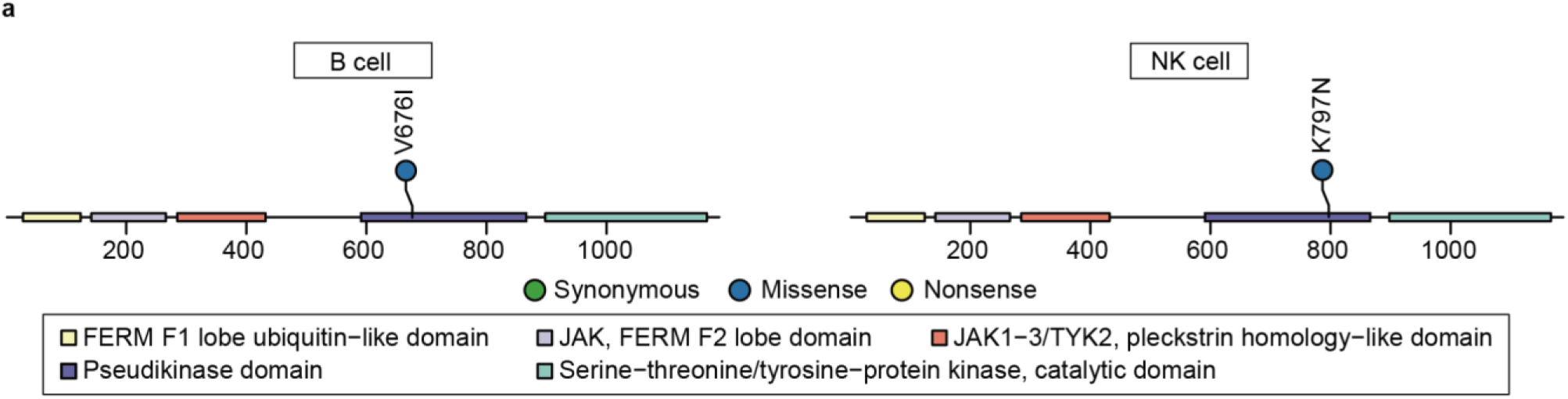
Lollipop plots of TYK2 mutations in TILs. a. Locations of TYK2 mutations in tumor-infiltrating B cells and tumor-infiltrating NK cells.

**Extended Data Figure 2.**
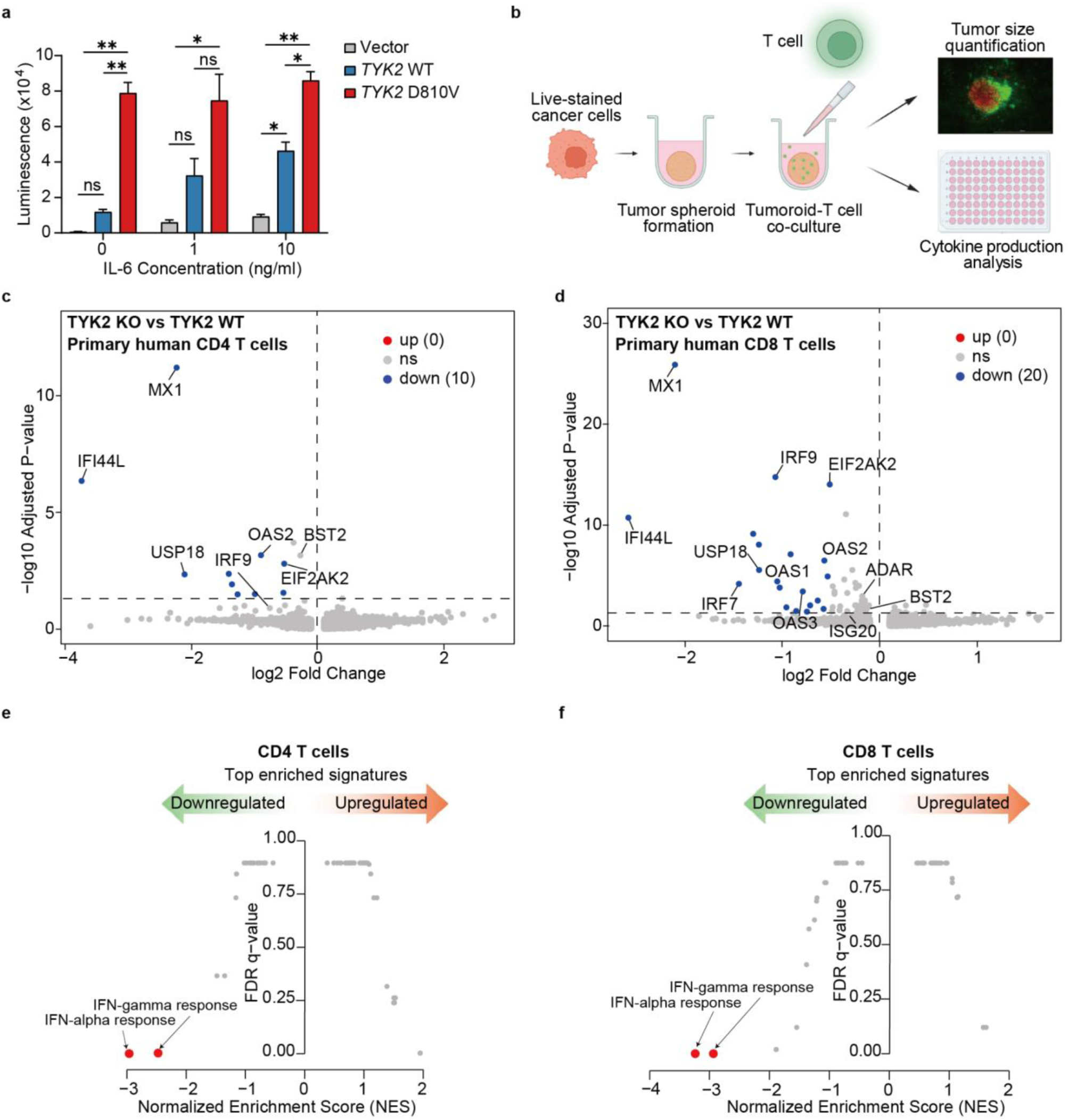
Functional characterization of *TYK2^D810V^* mutation and single-cell transcriptomic profiling of TYK2-deficient primary T cells. **a**. STAT3 reporter activity in TYK2-knockout 293T cells expressing empty vector, *TYK2^WT^* or *TYK2^D810V^*. b. A schematic of A549 tumor and T cell co-culture experiment. c. Volcano plot showing genes downregulated in TYK2-knockout primary human CD4⁺ T cells. d. Volcano plot showing genes downregulated in TYK2-knockout primary human CD8⁺ T cells. e. GSEA enrichment plot showing top enriched Hallmark gene sets ranked by NES in TYK2-knockout primary CD4⁺ T cells. f. GSEA enrichment plot showing top enriched Hallmark gene sets ranked by NES in TYK2-knockout primary CD8⁺ T cells.

**Extended Data Figure 3.**
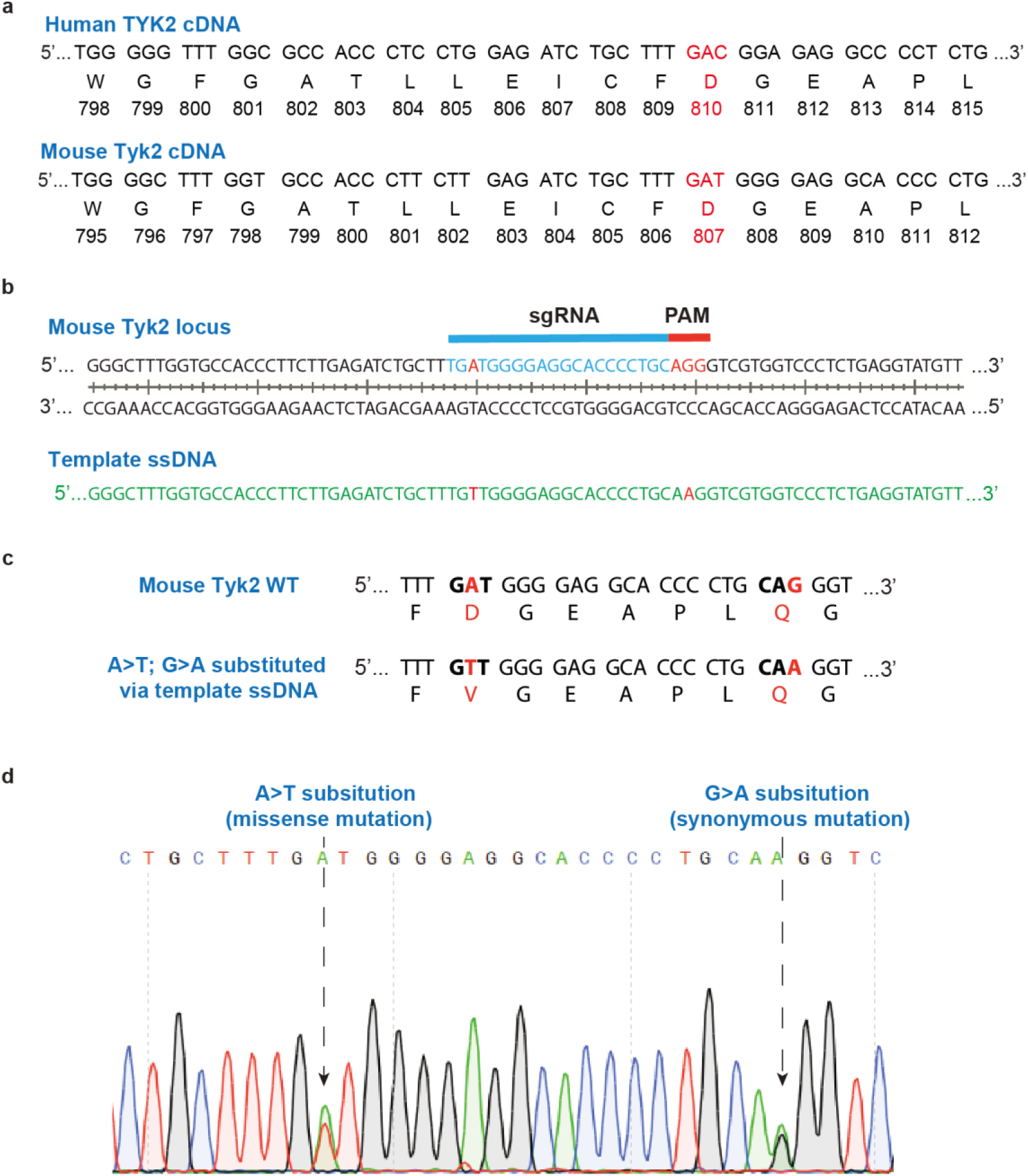
Design of *Tyk2^D807V^* mutant mice. **a.** Genomic sequences for human *TYK2^D810V^* and mouse *Tyk2^D807V^*. b. Single-stranded DNA (ssDNA) and sgRNA used with CAS9 to generate the *Tyk2^D807V^* mouse. c. Mouse TYK2 genomic sequences before and after CRISPR editing. **d.** Sanger sequencing for het *Tyk2^D807V^* point mutation mice.

**Extended Data Figure 4.**
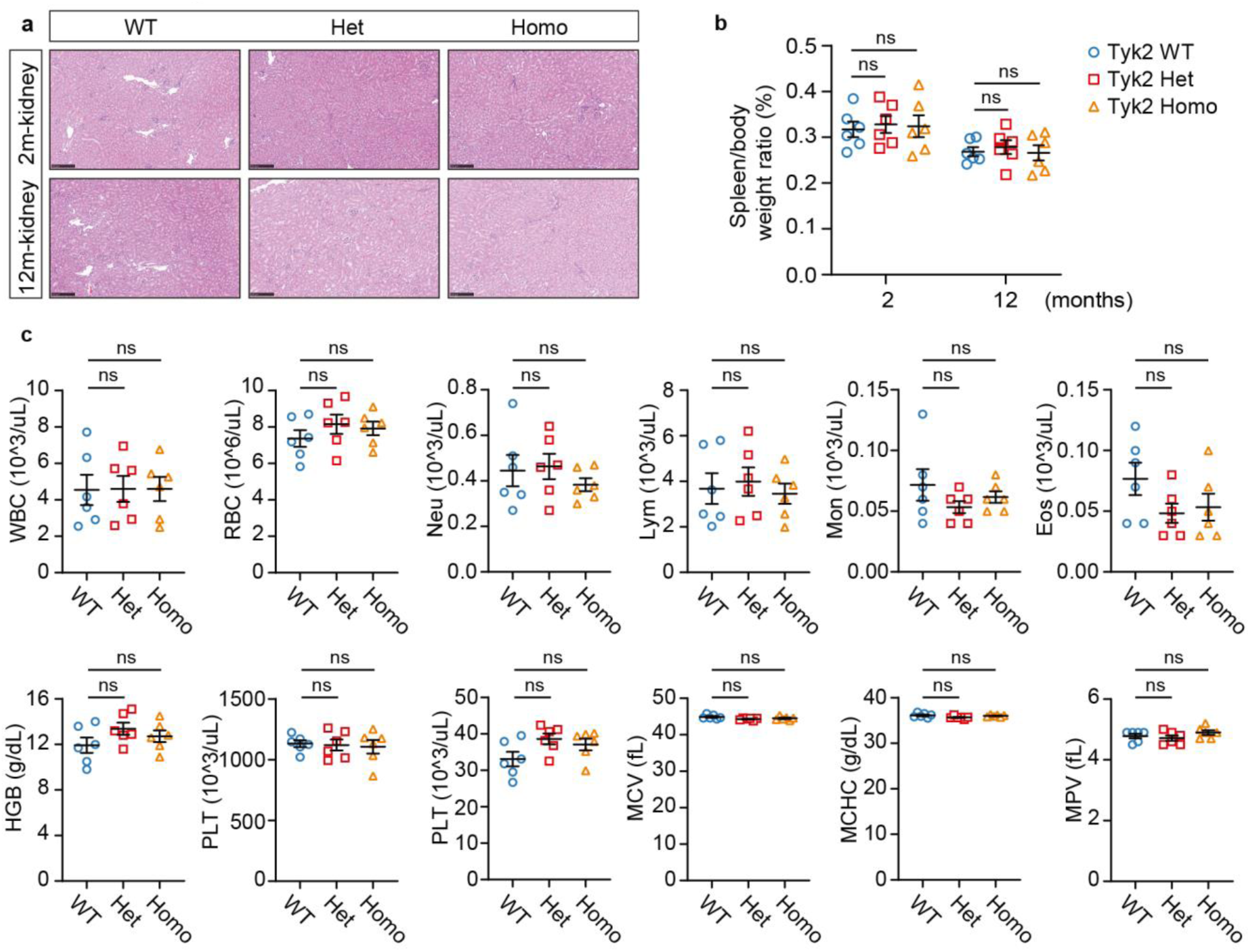
*Tyk2^D807V^* mutant mice appear healthy with no obvious signs of disease. a. H&E of kidney sections of 2- and 12-month-old het or homo *Tyk2^D807V^* mice and WT littermates (scale bars = 250 μm). b. Spleen-to-body weight ratios (n = 6 for each genotype). c. Complete blood counts of 2-month-old het or homo *Tyk2^D807V^* mice and WT littermates (n = 6 for each genotype). For data in this figure, mean ± s.e.m. is depicted. *P* values were determined by one-way ANOVA followed by Tukey’s multiple comparisons test. \**P* < 0.05; \*\**P* < 0.01; \*\*\**P* < 0.001; \*\*\*\**P* < 0.0001; ns, not significant.

**Extended Data Figure 5.**
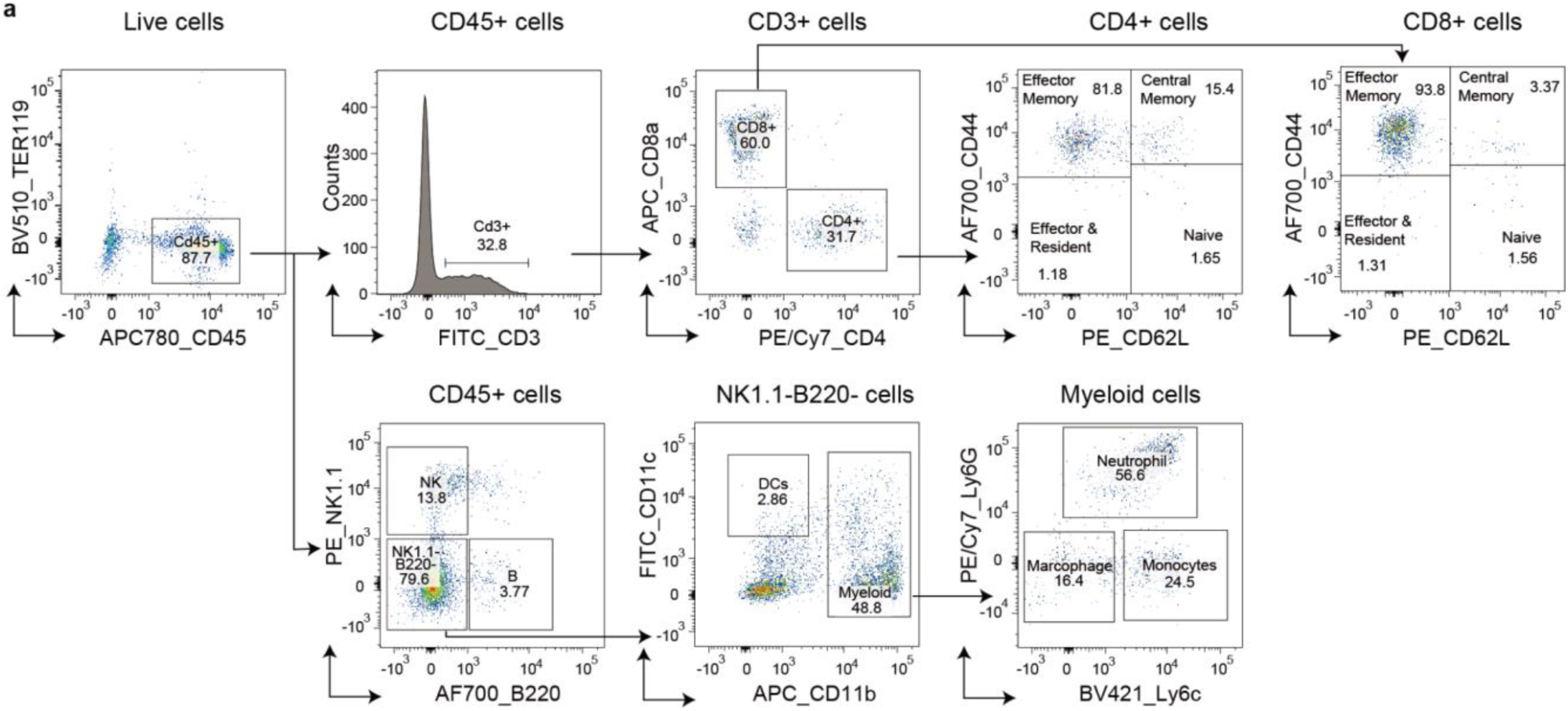
*Tyk2^D807V^* mutant mice showed increased T cell infiltration upon tumor challenge. a. Representative flow cytometry gating strategy for immune cell subset identification from tumor-infiltrating CD45⁺ cells.

**Extended Data Figure 6.**
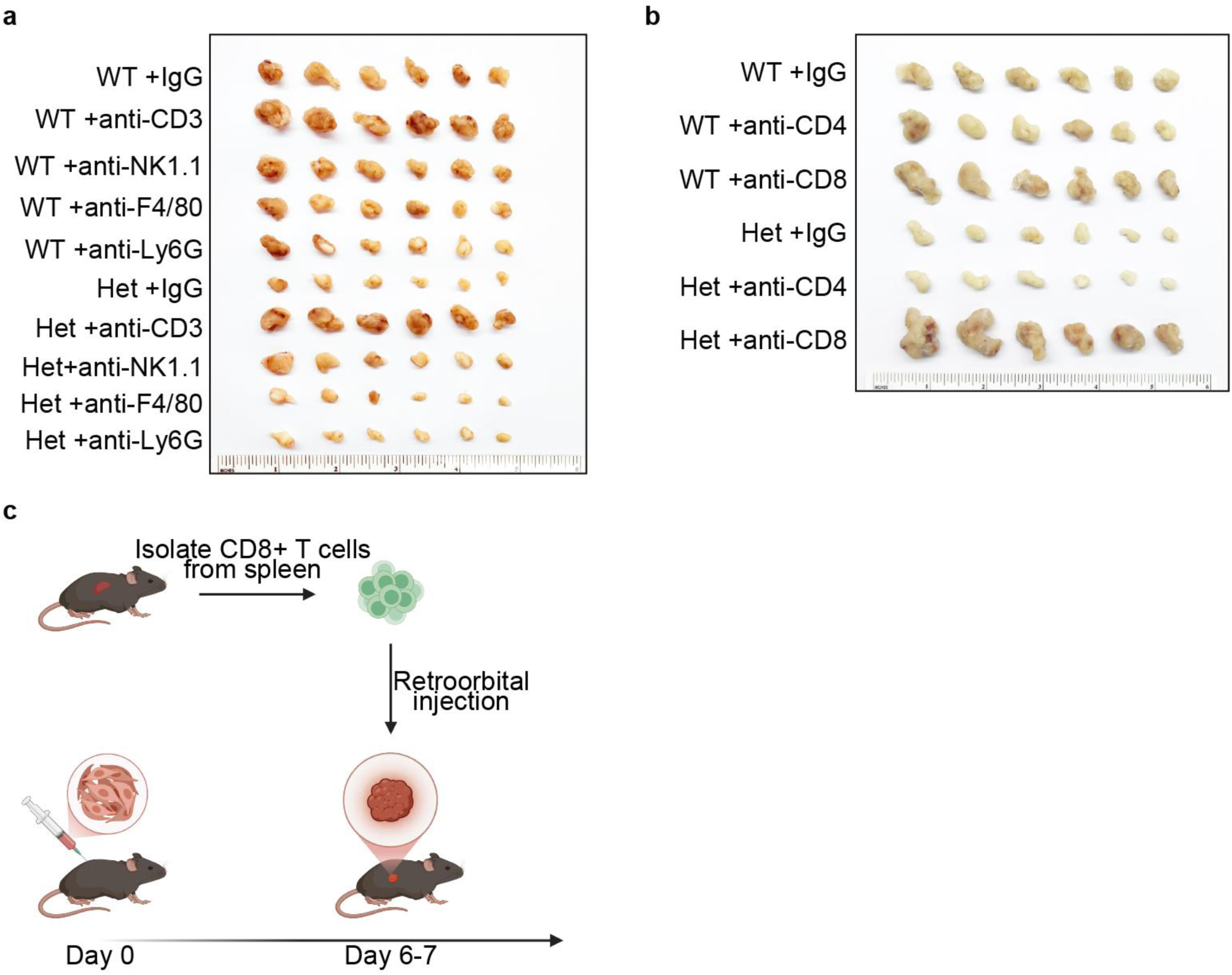
The anti-tumor effects of *TYK2^D807V^* mice are mediated by CD8⁺ T cells. a. MC38 syngeneic tumors from het *Tyk2^D807V^* and WT littermates treated with anti-IgG, anti-CD3ε, anti-NK1.1, anti-F4/80 and anti-Ly6G antibodies. Tumors from each group were ranked by size in descending order, and the top six are shown. b. MC38 syngeneic tumors from het *Tyk2^D807V^* and WT littermates treated with anti-IgG, anti-CD4 and anti-CD8β antibodies. Tumors from each group were ranked by size in descending order, and the top six are shown. c. Experimental schema for adoptive transplantation of CD8⁺ T cells.

